# Immortalization of human valve interstitial cells: an *in vitro* tool for studying fibro-calcific aortic valve degeneration

**DOI:** 10.1101/2024.09.26.615125

**Authors:** Donato Moschetta, Ilaria Massaiu, Francesca Bertolini, Vincenza Valerio, Valentina Rusconi, Donato De Giorgi, Veronika A. Myasoedova, Paolo Poggio

**Affiliations:** Centro Cardiologico Monzino IRCCS, 20138 Milan, Italy; Department of Biomedical, Surgical and Dental Sciences, University of Milan, Milan, Italy

## Abstract

Human primary valve interstitial cells (VIC) are essential for studying the detrimental fibro-calcification processes typical of aortic valve degeneration. However, their limited lifespan poses a significant challenge. To address this, we immortalized VICs to extend their lifespan and facilitate genetic manipulation, enabling the study of dysregulated molecular and cellular pathways in this pathology and the test of potential drugs *in vitro*. We achieved immortalization by transducing primary VICs with simian virus 40 gene and human telomerase large T gene. Post-immortalization, we conducted RNA sequencing on both primary and immortalized VICs to identify pathways regulation following infection. Additionally, we assessed whether immortalized VICs retained their mesenchymal markers and responded to the pro-osteogenic and pro-calcific stimuli similarly to primary VICs. Given the comparable data obtained from both cell types, we propose using immortalized VICs as a reliable tool for *in vitro* study of fibro-calcific aortic valve degeneration.

## Introduction

The use of primary cells has been indispensable for advancing scientific discoveries in cell biology. However, a significant limitation of these cells is their limited proliferative potential.^1^ Somatic cells become senescent after a finite number of divisions, a process triggered by various stressors such as irreparable DNA damage, telomere shortening, excessive oncogenic signals, non-genotoxic stress, and other unknown factors.^2^ Cellular senescence results in the arrest of cell proliferation, producing vital but non-mitotic cells with altered gene expression.^3^ These senescent cells are unsuitable for research as they do not respond to physiological, chemical, and physical stressors like normal cells and are not useful for *in vitro* pharmacological testing. Consequently, researchers must continuously isolate new primary cells from new donors, leading to significant variability.

Advancements in surgical techniques have led to less invasive procedures, complicating the acquisition of human specimens for primary cell isolation and this is particularly true for aortic valve replacement. Traditionally, surgical aortic valve replacement (AVR) provided ample aortic valve specimens for cell isolation. However, transcatheter aortic valve implantation (TAVI) is increasingly replacing AVR due to its benefits and less invasive nature. Since 2008, the percentage of patients undergoing TAVI has grown rapidly, while AVR procedures have decreased.^4^ Initially proposed for high-risk patients, TAVI is now also offered to medium and low-risk patients.^5^ This shift is expected to significantly reduce the availability of surgical AVR specimens and, consequently, human primary cells for research. Additionally, while animal models exist for studying aortic stenosis, none faithfully replicate the fibro-calcific processes observed in human diseased aortic valves.^6^ Hence, it is imperative to explore new strategies to study aortic valve stenosis.

Valve interstitial cells (VIC) are considered a reliable tool for investigating the molecular and cellular mechanisms of fibro-calcific aortic valve degeneration (FCAVD).^7^ Immortalizing these cells through genetic manipulation can address the issues of biodiversity and cell availability for *in vitro* experiments. Typically, immortalization involves p53-mediated terminal proliferation abrogation^8^ and/or activation of telomerase reverse transcriptase (hTERT), which prevents senescence due to telomere shortening.^9,10^ In this study, we immortalized human primary VICs isolated from aortic valve leaflets by genetically activating the simian virus 40 (SV40) and hTERT genes, and subsequently evaluate their characteristics.

## Results

### Transcriptome analysis of immortalized valve interstitial cells

We consecutively infected VICs isolated from FCAVD patients with simian virus 40 (SV40) and human telomerase T (hTERT) under a cytomegalovirus promoter to ensure their constitutive transcription. The immortalized VICs (iVICs) demonstrated the ability to be expanded *in vitro* for at least 30 passages, in contrast to primary VICs, which could only be cultured for 5-6 passages before reaching replicative senescence.

To assess the impact of immortalization, we performed poly-adenylated RNA transcriptome sequencing on cells from the same donors (n = 7), both pre- and post-immortalization. Multidimensional scaling analysis revealed distinct clustering between primary and immortalized VICs, indicating significant transcriptional changes (Figure 1A). Differential gene expression analysis identified over 1,000 genes with significant enrichment in either VICs or iVICs [log2(fold change) > |±1|; adjusted p < 0.01; Figure 1B and Supplemental Table S1]. Functional analysis of these gene expression profiles highlighted that pathways related to mitotic division, such as chromosome segregation, DNA replication, organelle fission, and cell cycle, were positively regulated in iVICs (Figure 1C). Conversely, pathways enriched in primary VICs included those involved in generation of precursor metabolites and energy, intracellular trafficking, and extracellular matrix organization, reflecting the characteristics of non-replicating cells in the final stages of differentiation. Thus, our findings demonstrate that immortalized VICs retain key functional characteristics of primary VICs while exhibiting enhanced proliferative capacity.

**Figure 1.**
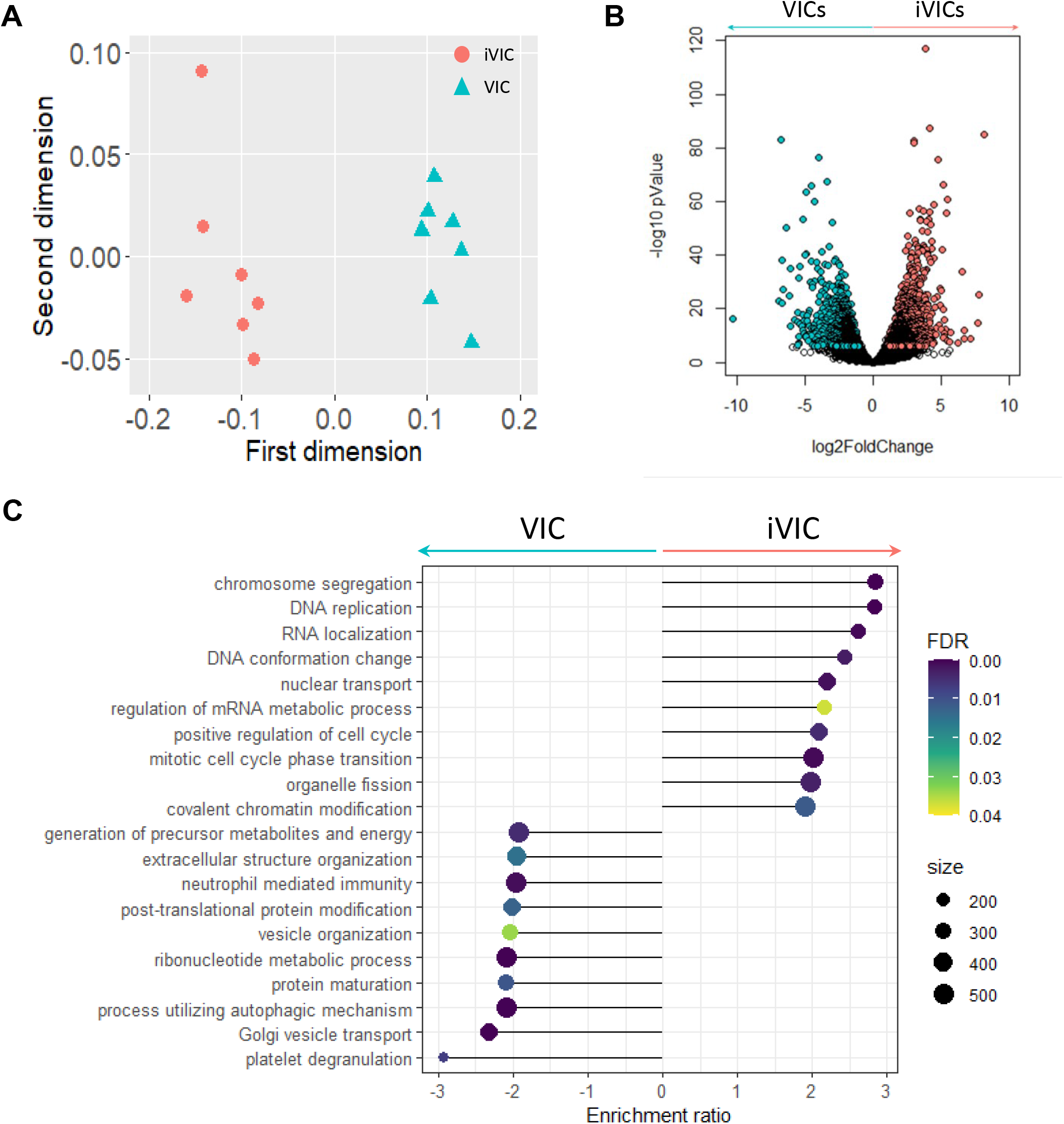
Transcription sequencing. Principal component analysis (**A**) and volcano plot of differentially expressed genes (**B**) coming from transcription sequencing of valve interstitial cells (VIC) and immortalized VICs (iVIC). Light red and light blue dots represent upregulated genes in iVICs and VIC, respectively, with both the adjusted pValue < 0.01 and the absolute log2 fold change (logFC) > 1 (**C**) Functional analysis of molecular and cellular processes activated/repressed by immortalization. Light blue: VIC; light red (iVIC).

### Immortalized valve interstitial cells characterization

The proliferative capacity of VICs and iVICs was assessed using the Ki67 proliferation marker. Both VICs, at low passage (lower than 5^th^ passage), and iVICs, at high passage (higher than 15^th^ passage), exhibited similar Ki67 expression levels (39.98 ± 7.26% vs. 40.67 ± 5.84%, respectively; p = 0.83; Figure 2A and 2B), indicating that iVICs retain their proliferative potential even at higher passages, unlike VICs which lose this ability. To confirm the mesenchymal phenotype of iVICs, we evaluated the expression of vimentin (VIM) and cluster of differentiation 90 (CD90) as previously tested in primary VICs.^11^ As expected, all iVICs co-expressed these markers, confirming their mesenchymal identity (Figure 2C and 2D). Furthermore, the calcification potential of VICs and iVICs was quantified after seven days in pro-osteogenic (PO) and pro-calcific (PC) media. Both cell types exhibited proportional calcification in response to PO and PC media. Primary VICs showed significant increases in calcification (+4.67±4.27 and +12.16±14.62, respectively; both p < 0.01; Figure 2E) compared to untreated cells, while iVICs demonstrated similar trends (+1.37±0.36 and +12.22±11.55, respectively; both p < 0.01; Figure 2F). Thus, these results corroborate the use of iVICs as a robust *in vitro* model for studying FCAVD.

**Figure 2.**
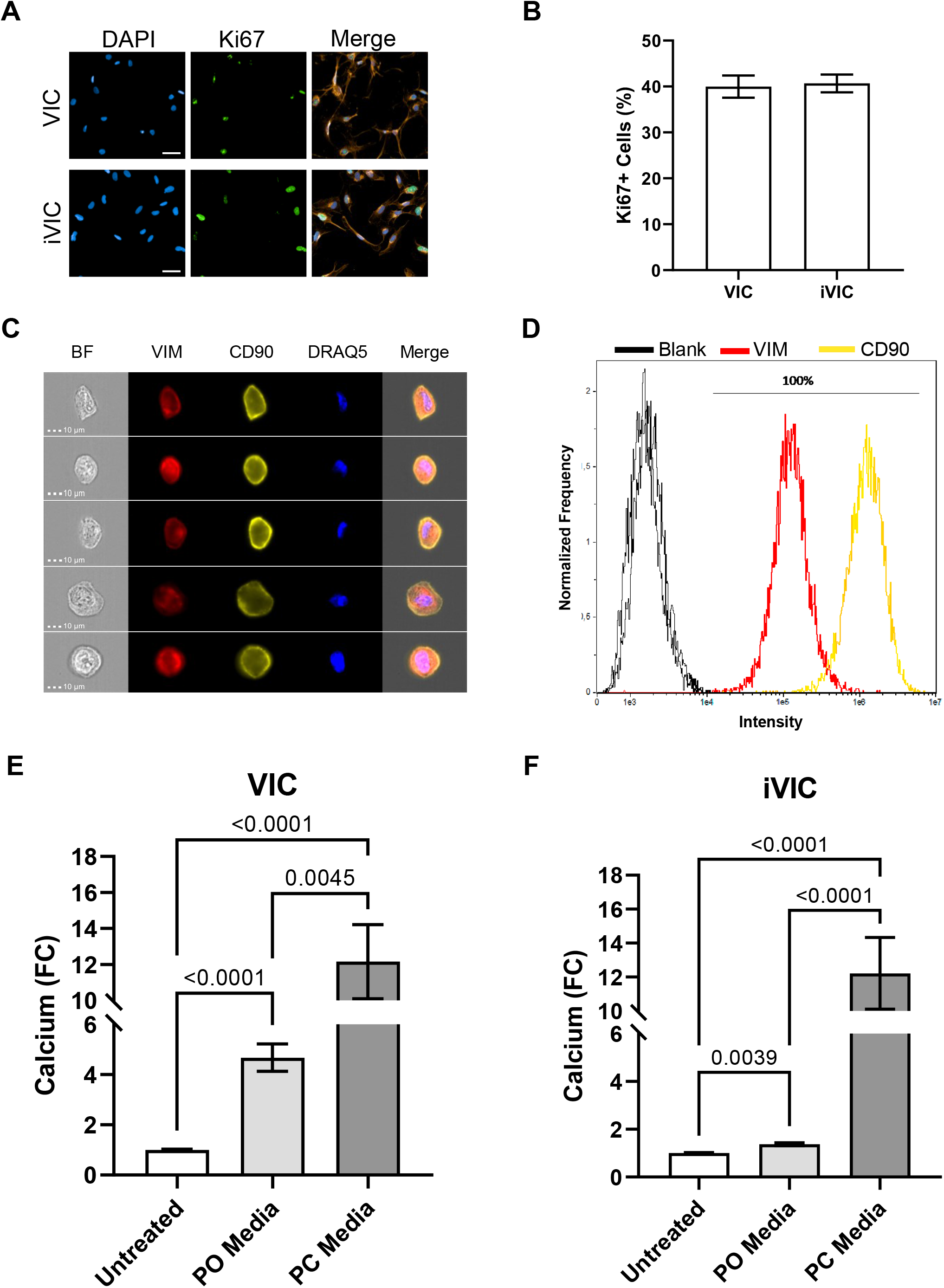
Immortalized VIC characterization. Representative images (**A**) and relative quantification (**B**) of ki67 (proliferation marker) in VICs and iVICs. Representative images (**C**) and relative quantification (**D**) of iVICs stained for vimentin (in red; Vim) and CD90 (in yellow) acquired by ImageStreamX. Extracellular calcium quantification in VIC (**E**) and iVIC (**F**) after treatment with pro-osteogenic (PO) and pro-calcific (PC) media. Data are showed as mean ± standard error of mean (SEM) and expressed ad fold change (FC) versus untreated. ****: p < 0.0001.

## Discussion

FCAVD progression is characterized by the activation of several cellular and molecular processes.^7^ The most detrimental ones are caused by VICs activation and the consecutive pathological rearrangement of extracellular matrix, which bring to the accumulation of fibrotic fibers and calcium deposition.^12^ Most of the research studies on molecular processes activated during FCAVD have been performed on primary VICs *in vitro*, due to the lack of animal model that fully recapitulate the pathology.^6^ Indeed, mice and rats have a different aortic valve layer ultrastructure, characterized by the absence of the typical human three-layers structure.^13^ For this reason, even if they evolve valvular calcification after genetic or pharmacological manipulation, they do not show hemodynamic changes unless fed with high cholesterol Western diet and aged for 6-12 months.^6^ The only animal model that spontaneously evolves FCAVD and presents most of the common features showed in human disease is the pig.^6,14^ However, the use of pigs as an experimental model is reduced due to the time and cost involved in experimenting with large animals. Therefore, the limitation of current animal models and the central role of VICs in the onset and progression of FCAVD have made the latter the experimental tool of choice to study the pathology and screen for pharmacological intervention.^15^ In addition, thanks to the advancement of non-invasive surgery, more and more patients undergo TAVI instead of AVR, thus leaving less aortic valve leaflets available for primary VIC isolation. Given the urgent need to identify a molecular target for FCAVD, coupled with the limited proliferative capacity of primary VICs and significant biological variability, it is imperative to develop a new cellular model to study the disease.

In pursuit of this goal, we embarked on the genetic manipulation of primary VIC, bulding up on previously validated protocols.^16,17^ By transducing the cells with lentiviral particles carrying the SV40 and hTERT genes, we enabled them to circumvent the DNA repair mechanisms activated by p53 and to replicate indefinitely without telomere shortening. Ki67 staining, a renowned proliferation marker,^18^ confirmed that iVICs were actively replicating even at the 30^th^ passage, whereas primary VICs could not be used beyond the 6^th^ passage. This was further substantiated by differential gene expression and functional analyses, which revealed that the genes differentially expressed in iVICs were predominantly associated with the cell cycle and the evasion of the DNA damage checkpoints. This transformation significantly enhanced cell proliferation without impacting other cellular pathways.

Although immortalization accelerates the cell proliferation rate and increases the number of population doublings, it does not alter the differentiation status of our cells. Considering the iVIC characterization, they consistently expressed vimentin and CD90, mesenchymal markers typical of VICs,^11,19^ confirming that they did not de-differentiate. Finally, we assessed the calcification potential of iVICs by exposing them to pro-osteogenic and pro-calcific media, and found that they calcify to the same extent as primary VICs.

Collectively, these data confirmed the potential use of iVICs as experimental tool for *in vitro* studies of the molecular pathways activated in VICs as well as to screen for new pharmacological compounds.

## Methods

### Valve interstitial cell isolation

Aortic valve interstitial cells (VICs) were isolated by aortic valve leaflets within 30 minutes by the surgical resection. Each leaflet was incubated for 20 min at 37 °C in 2 mg/mL type II collagenase (Worthington Biochemical Corp., Lakewood, NJ, USA) in cell culture medium (advanced Dulbecco’s modified Eagle’s medium (Ad DMEM, Life Technologies, Carlsbad, CA, USA) containing 10% fetal bovine serum (FBS, Microtech, Naples, Italy), 1% penicillin (Life Technologies, Carlsbad, CA, USA), 1% streptomycin (Life Technologies, Carlsbad, CA, USA), and 1% l-glutamine (Life Technologies, Carlsbad, CA, USA)). This step was implied to remove the endothelial cells. Then, leaflets were moved away in new fresh medium and finely shredded with a scalpel, before incubating overnight in 2 mg/mL type II collagenase in cell culture medium at 37 °C to process the extracellular matrix proteins. At the end of incubation, the resulting valve interstitial cells (VIC) were plated in tissue culture plates in complete Ad DMEM and maintained at 37°C and 5% CO2. All the experiments were performed on cultured cells between the second and fifth passage.

### Lentiviral transduction of VIC

Primary VICs were immortalized by transduction with commercial lentiviral particles (GeneCopoeia) carrying out the encoding sequences for SV40 large T antigen (LP724-100; GeneCopoeia, Rockville, MD, USA) and human telomerase reverse transcriptase (hTERT) enzyme (LP729-100; GeneCopoeia, Rockville, MD, USA) under cytomegalovirus promoter. VICs were transduced as follow: 2.0 × 104 cells were plated in twenty-four-well tissue culture plates and incubated at 37°C, 5% CO2 until they reached up to 80% confluence. Afterwards, medium was changed with fresh complete Ad DMEM with 8 μg/ml of polybrene and SV40 large T lentiviral particles 10 molecules of infection (MOI). Cells were then incubated overnight at 37°C, 5% CO2, for 48 hours, before changing the medium. Cells were left in culture until they reach the confluence, then splitted 1 to 5. After two passages, SV40-immortalized VICs were plated at the same confluence and treated in the same way adding hTERT lentiviral particles. When confluent, double immortalized cells were selected by splitting 1 to 5 until at least 30 passages.

### RNA extraction and sequencing

RNA extraction was performed from VICs and iVICs using Total RNA Purification Plus Kit (Norgen Biotek Corp., Thorold, ON, Canada) according to the manufacurer’s instructions. RNA quantification was determined with a spectrophotometer Infinite® 200pro (TECAN) spectrophotometer. We prepared cDNA libraries following the recommendations of the Nanopore cDNA-Seq protocol for the SQK-PCS109 kit, as previously done in our lab.^20^ Briefly, we employed RT primers to convert only poly-adenylated RNA into cDNA. For the multiplex run, we used six different ad hoc designed barcoded sequences. cDNA synthesis was performed using 50 ng of total RNA per sample. RT and strand-switching primers were provided by ONT with the SQK-PCS109 kit. Following RT, PCR amplification was performed using the LongAmp Taq 2X Master Mix (New England Biolabs, Ipswich, MA, USA) and the following cycling conditions: 1 cycle (95 °C for 30 s), 18 cycles (95 °C for 15 s, 62 °C for 15 s, and 65 °C for 3 min), and 1 last cycle (65 °C for 15 min). PCR products were purified using Agencourt AMPure XP beads (Beckman Coulter, Brea, CA, USA). The cDNA sequencing libraries were prepared using a total of 200 fmol of cDNA each. Nanopore libraries were sequenced using a MinION Mk1B sequencing device with R9.4 flow cells. Sequencing was controlled and data were generated using ONT MinKNOW software (v3.4.12). Runs were terminated after 48 h and FAST5 files were generated.

### Data processing, gene expression and functional analysis

DNA bases were called from FAST5 files using ONT Guppy GPU (v5.0.15) in high accuracy mode.^21^ Reads with an average Phred quality score, which measures the confidence based on the estimated error rate, lower than 7 were discarded. Reads were aligned to the 22 diploid chromosomes of the GRCh38 human genome reference with minimap2 (v2.1, default parameters except for -ax splice).^22^ SAM-to-BAM format conversion as well as an assessment of the alignment quality were performed using Samtools (v 1.10).^23^ The FeatureCounts software (v2.0.0),^24^ included in the Subread package, was used to count the mapped reads.

The raw counts were processed using the R software environment (v 3.6.0). In particular, the genes were annotated based on the ENSEMBL ID and log2 counts per million mapped reads (CPM) were computed).^25^ For the multiplex run, the quality-checked reads were de-multiplexed and trimmed for barcodes using the Cutadapt function (v1.15),^26^ before the alignment and counting procedures. Genes with a read count greater than 3 were deemed as expressed.

Differential expression analysis between immortalized and primary samples was performed through the limma R/Bioconductor package.^27^ The genes with absolute value of log2 fold-change (|logFC|) > 1 and FDR-adjusted pValue < 0.05 were considered as differently expressed. The robustness of the differential expression analysis results was assessed by exploring the histogram of the pValue distribution.

Gene set enrichment analysis (GSEA) was conducted using GSEA pre-ranked tool (v 4.0.3) (17644558) and the biological functions were inferred by taking advantage of gene ontology biological processes (GO-BP), Reactome, WikiPathways, and BioCarta databases. To reduce redundancy and visually interpret GSEA results, the most significant GO-BP/Reactome/BioCarta pathway gene sets were identified by a false discovery rate (FDR) < 0.1. The network was drawn through the Enrichment Map software (v.3.3.2)^28^ implemented as a plug-in in Cytoscape (v.3.8.2).^29^

### Imaging Flow Cytometry Analyses

VIC characterization was performed by an ImageStream X flow cytometer, which combines flow cytometry with microscopy technology (ImageStream X Mark II, Amnis). VICs were detached through TripLE Select (Gibco, Carlsbad, CA, USA), fixed cells in paraformaldehyde, incubated in NH4Cl 50 mM to break down cell auto-fluorescence, permeabilized in saponin 0.2%, and resuspended in 100 μL of PBS with 5 mM ethylenediaminetetraacetic acid (EDTA), 1% bovine serum albumin (BSA), and 0.2% saponin. Then, cell samples were incubated for 30 min at room temperature using specific conjugated-antibodies against CD90–phicoeritrin (Becton Dickinson PharmingenTM, Franklin Lakes, NJ, USA), vimentin–AlexaFluor 488 (Becton Dickinson PharmingenTM, Franklin Lakes, NJ, USA), and DRAQ5 (eBioscience, St. Clara, CA, USA), a nuclear stain. Samples were then washed with FACS buffer (PBS containing 5 mM EDTA and 1% BSA) by centrifugation for 5 min at 600× g. Finally, cells were resuspended in FACS buffer and analyzed. A total of 30,000 events were acquired in the single cells and focused gated area. Image analysis was performed using the IDEAS 6.2 software.

### Immunofluorescence

VIC and iVICs have been seeded (5000 cells/well) in Advance DMEM medium with 10% of FBS. The day after, cells have been washed and fixed in paraformaldehyde 4%. Then, cells have been gently washed with PBS and incubated with of wheat germ agglutinin (WGA; Thermo Fisher Scientific, Waltham, MA, USA). Then we permeabilized cells with cold methanol and performed the blocking in 5% BSA, 0.5% donkey serum and 0.2% tritonX100 PBS. Then we stained overnight with the anti-Ki67 (Cell Signaling Technology, Boston, MA, USA) and the day after with anti-mouse conjugated with alexa fluor 488 Jackson Immunoresearch, Cambridge, UK) for two hours at room temperature. After wash the secondary antibody in PBS, we put a drop of vectashield containing DAPI and acquire images with Operetta CLS.

### Calcium assay

Thirty thousand of cells were seeded in a 24 well plate (Merck, Darmstadt, Germany) in cell culture medium. After two days (when the confluence was about 80%) media was changed and the treatments were added. We use pro-osteogenic medium (β-glycerophosphate 10 mM and ascorbic acid 50 µg/mL; βGAA) and pro-calcific medium (inorganic phosphate 2 mM (Pi)). Treatments and media were changed every other day for seven or fourteen days.

After removing the culture media, extracellular calcium crystals were dissolved with 0.6 M hydrochloric acid (HCl) for 5 h in gentle agitation. Subsequently, HCL samples were recovered and stored +4°C; meanwhile, the layer of cells present in each well was treated with 0.1 M sodium hydroxide (NaOH) and 0.1% sodium dodecyl sulfate (SDS) to ensure cellular lysis necessary for total protein quantification. Extracellular calcium quantification was performed using a colorimetric assay kit (BioVision Inc., Milpitas, CA, USA) following the manufacturing company’s protocol. The absorbance was read off at the Infinite® 200pro (TECAN) spectrophotometer. Raw data on calcium concentration were normalized on the total protein measured by the bicinchoninic acid (BCA) protein assay kit (Thermo Fisher Scientific, Waltham, MA, USA).

## Author contributions statement

D.M. and P.P. conceived the study and the experimental plan. D.M., F.B., D.D.G., V.V., and V.R. conducted all the experiments. D.M. and I.M. analyzed the results. V.A.M. enrolled the subjects for the study. D.M. and P.P. write the original manuscript. All authors reviewed the manuscript.

## Notes

### Competing Interest Statement

The authors have declared no competing interest.

